# Shorter telomere length is associated with a more recent diagnosis of coeliac disease

**DOI:** 10.1101/331405

**Authors:** Pierre Goorkiz, Nerissa L Hearn, Saskia van der Kooi, Christine L Chiu, Joanne M Lind

**Affiliations:** School of Medicine, Western Sydney University, NSW, Australia.; Faculty of Medicine and Health Sciences, Macquarie University, NSW, Australia

## Abstract

**Background:** Coeliac disease (CD) is an autoimmune disease that causes an inappropriate inflammatory immune response to dietary gluten. Telomere length is a marker of biological ageing and is reduced in several autoimmune conditions. This observational study measured salivary telomere length (TL) in gluten-free diet (GFD) treated CD individuals to determine if CD, and length of time on a GFD, is associated with salivary TL.

**Methods:** Clinical and demographic information was collected from CD individuals currently treated with a GFD and healthy non-affected controls. Only participants aged under 35 years at recruitment were included. Relative telomere length was measured using quantitative PCR in oral mucosa collected from saliva. Linear regression was used to determine whether salivary TL was associated with CD, or length of time on a GFD, adjusting for age and sex.

**Results:** This study included 79 participants, 52 GFD-treated CD and 27 non-affected controls. No significant difference in salivary TL between individuals with treated CD and controls was found. Within CD individuals, salivary TL was associated with length of time on a GFD, with individuals who started a GFD ≤3 years ago having shorter salivary TL compared to those who started a GFD > 3 years ago (0.37±0.05 vs 0.50±0.04; p=0.002).

**Conclusion:** Our findings indicate that salivary TL shorten while CD is untreated, however following treatment on a GFD, they appear to recover to those seen in unaffected controls. This highlights the importance of early diagnosis and initiation of GFD to minimise mucosal damage and telomere shortening, to enable TL to recover.

## Introduction

Coeliac disease (CD) is an autoimmune disorder characterised by damage to the small intestine in genetically predisposed individuals [1]. In individuals with CD, exposure to dietary gluten triggers an immune response producing oxidative stress, inflammation and histological changes within the intestinal mucosa [2-4]. The only treatment is a lifelong gluten free diet (GFD) which, results in remission of immune mediated intestinal damage and symptom resolution [4].

Telomeres are repetitive DNA sequences that cap the end of chromosomes, protecting them from degradation and chromosome fusion[5]. These structures progressively shorten with each cell division, until they reach a critical length and induce cell senescence and/or cell death. Telomere length (TL) is therefore widely considered as a marker of biological ageing. Shorter telomeres are associated with states of inflammation and oxidative stress, where progressive cell damage results in greater cell turnover, and free radical mediated damage results in ‘clipping’ of telomere DNA during each cycle of division [6-8]. The effects of inflammation and oxidative stress on TL have been shown to be reversible, with length gradually recovering when these stressors are removed [9].

Shorter TL have been associated with autoimmune conditions, asthma, and depression [10-12]. In CD, shorter TL have been reported in small intestinal biopsy samples and peripheral blood lymphocytes of individuals with CD compared to healthy controls [13]. However, these studies measured TL in individuals with active or untreated CD, that is, in individuals consuming a gluten-containing diet. No studies have investigated whether TL can recover after treatment of CD with a GFD. The aim of this study was to determine whether salivary TL differed between CD individuals currently on a GFD and age-matched healthy controls in a population of individuals under 35 years of age. This study also aimed to determine whether the length of time on a GFD was associated with TL recovery in individuals with CD.

## Methods

### Participant recruitment and inclusion criteria

Recruitment was carried out between April 2014 and December 2016. Individuals were recruited from the gastroenterology clinic at Campbelltown Hospital, NSW, and from the general population at the 2014, 2015 and 2016 Gluten-Free Expos in Sydney and Melbourne, Australia. Following written informed consent, individuals were asked questions regarding their health and disease status, as previously described [14]. Saliva samples were collected from all participants using the Oragene DNA OG500 self-collection kits (DNA Genotek, Canada). This study was approved by the Western Sydney University Research Ethics Committee (approval number H10513) and was carried out in accordance with the ethical standards laid down in the 1964 Declaration of Helsinki and its later amendments.

Only individuals between the age of 18 and 35 at the time of recruitment were included in the study. Individuals were defined as having CD if they fulfilled the following criteria: had been diagnosed with CD via duodenal biopsy by a gastrointestinal specialist; were currently on a GFD; and carried at least one HLA-DQ2 or HLA-DQ8 haplotype. Endoscopy reports were obtained for a subset of CD participants to verify CD histology and confirm their CD status. Individuals were classified as non-affected controls if they reported not having CD or CD associated symptoms and were not on a GFD.

BMI was analysed as a categorical variable according to World Health Organization guidelines [15]. Individuals were classified as current, past, or non-smokers, and alcohol consumption was categorised into zero, 1-2, and 3-7 standard drinks per week. The length of time on a GFD was calculated by subtracting the date of commencement of a GFD from the recruitment date. CD individuals were dichotomised ≤3 years since starting a GFD and >3 years since starting a GFD. Participants reported if they had ever been diagnosed with cancer, depression, asthma, or any of the following autoimmune conditions: Type 1 diabetes mellitus; Autoimmune thyroid disease; Rheumatoid arthritis; Lupus; Dermatitis herpetiformis; or Psoriasis. Data from depression, asthma and autoimmune condition variables were combined to generate the variable ‘associated conditions’ as the prevalence of each individual condition was low. Individuals with missing data; a history of cancer; or who were current or past smokers, were excluded.

### DNA extraction and HLA genotyping

Collected saliva was stored at room temperature until DNA extraction, as per the manufacturer’s recommendations. Genomic DNA was extracted as per the Oragene prep IT L2P (DNA Genotek, Canada) protocol, purified using the Qiagen DNA mini kit (Qiagen, Germany) spin column protocol, and the samples were stored at −20°C until analysis. All samples were genotyped for the CD susceptibility haplotypes HLA-DQ2 and HLA-DQ8 using TaqMan SNP genotyping assays as previously described[16].

### Measurement of relative telomere lengths

Relative telomere length in saliva was measured using quantitative PCR as previously described[17, 18]. This method expresses telomere length as a ratio (T/S) of telomere repeat copy number (T) to haemoglobin subunit beta (*HBB*) single copy gene (S) within each sample. Therefore, a higher T/S corresponds to longer telomeres. All samples were run in triplicate, and a calibrator was included on each plate, and the coefficient of variability (CV) between plates was 8.44%. Cycle threshold (Ct) values for each sample were calculated using the MxPro QPCR software (Stratagene, Agilent Technologies, USA). Triplicate Ct values were averaged and the quantity of each sample was calculated relative to the calibrator using the delta-delta Ct method [19].

### Statistical analysis

Prior to analysis, T/S ratio values were transformed into natural logarithms to obtain a Gaussian distribution. Multinomial logistic regression was used to determine if there were significant differences in demographic and lifestyle factors between the GFD ≤3yrs CD individuals and unaffected controls, and the GFD >3yrs CD individuals and unaffected controls. Generalised linear models (GLZM) was used to compare salivary TL between the CD groups and unaffected controls, and between the GFD ≤3yrs CD and GFD >3yrs CD, adjusting for age. P values <0.05 were considered statistically significant. All analysis was performed using the IBM SPSS statistics software (version 24).

## Results and discussion

A total of 79 individuals, 52 CD and 27 non-affected controls were included in the study (Fig 1). These individuals were aged between 18 and 35 years, had not been previously diagnosed with any type of cancer, and reported having never smoked. There were significantly more female participants compared to males (88.6% vs 11.4%). There were no significant differences in age, BMI, alcohol consumption or presence of associated conditions between individuals with CD and non-affected controls (Table 1). Age, sex, BMI, alcohol consumption, or presence of associated conditions were not associated with salivary TL, however as age is a known predictor of TL, all subsequent analyses were adjusted for age. Stratifying the CD cohort by length of time on a GFD found CD individuals who had commenced a GFD ≤3 years ago had shorter TL when compared to non-affected controls, but this was not significant (0.37±0.05 vs 0.44±0.04; p=0.12). While for CD individuals who had commenced a GFD >3 years ago, no difference in TL was observed when compared to non-affected controls (0.50±0.04 vs 0.44±0.04; p=0.34). Within the CD cohort, length of time on a GFD was significantly associated with relative salivary TL, with CD individuals who had commenced a GFD ≤ 3 years ago having shorter TL compared with individuals who had removed gluten from their diet > 3 years ago (0.37±0.05 vs 0.50±0.04; p=0.002) (Fig 2).

**Table 1.** Demographic and lifestyle factors of individuals with coeliac disease compared with controls

| Characteristic |  | <u>Control</u> | <u>CD ≤3yrs</u> |  | <u>CD >3yrs</u> |  |
| --- | --- | --- | --- | --- | --- | --- |
|  |  | n=27(%) | n=20 (%) | OR [95% CI]; p-value | n=32 (%) | OR [95% CI]; p-value |
| Age (mean±SE years) |  | 22.9 ± 0.8 | 24.0 ± 1.0 | 0.07 ± 0.08; p=0.36* | 25.4 ± 0.7 | 0.15 ± 0.07; p=0.03* |
| Sex | Male | 5 (18.5) | 2 (10.0) | 1.00 | 2 (6.2) | 1.00 |
|  | Female | 22 (81.5) | 18 (90.0) | 0.49[0.09 – 2.82]; p=0.42 | 30 (93.8) | 0.29[0.05 – 1.65]; p=0.17 |
| BMI (kg/m<sup>2</sup>) | <25 | 21 (77.8) | 14 (70.0) | 1.00 | 23 (71.9) | 1.00 |
|  | 25-30 | 5 (18.5) | 4 (20.0) | 1.20[0.27 – 5.26]; p=0.81 | 5 (15.6) | 0.91[0.23 – 3.61]; p=0.90 |
|  | 30+ | 1 (3.7) | 2 (10.0) | 3.00[0.25 – 36.33]; p=0.39 | 4 (12.5) | 3.65[0.38 – 35.34]; p=0.26 |
| Alcohol (drinks/week) | 0 | 14 (51.9) | 12 (60.0) | 1.00 | 13 (40.6) | 1.00 |
|  | 1-2 | 10 (37.0) | 6 (30.0) | 0.70[0.20 – 2.50]; p=0.58 | 11 (34.4) | 1.19[0.38 – 3.70]; p=0.77 |
|  | 3-7 | 3 (11.1) | 2 (10.0) | 0.79[0.11 – 5.46]; p=0.80 | 8 (25.0) | 2.87[0.62 – 13.22]; p=0.18 |
| Associated Conditions | No | 16 (59.3) | 11 (55.0) | 1.00 | 18 (56.3) | 1.00 |
|  | Yes | 11 (40.7) | 9 (45.0) | 1.19[0.37 – 3.83]; p=0.77 | 14 (43.8) | 1.13[0.40 – 3.19]; p=0.82 |

CD = coeliac disease; OR = Odds Ratio; CI = Confidence Interval; BMI = body mass index; SE = standard error.
\*Linear variable, therefore β±SE is reported.

**Figure 1.**
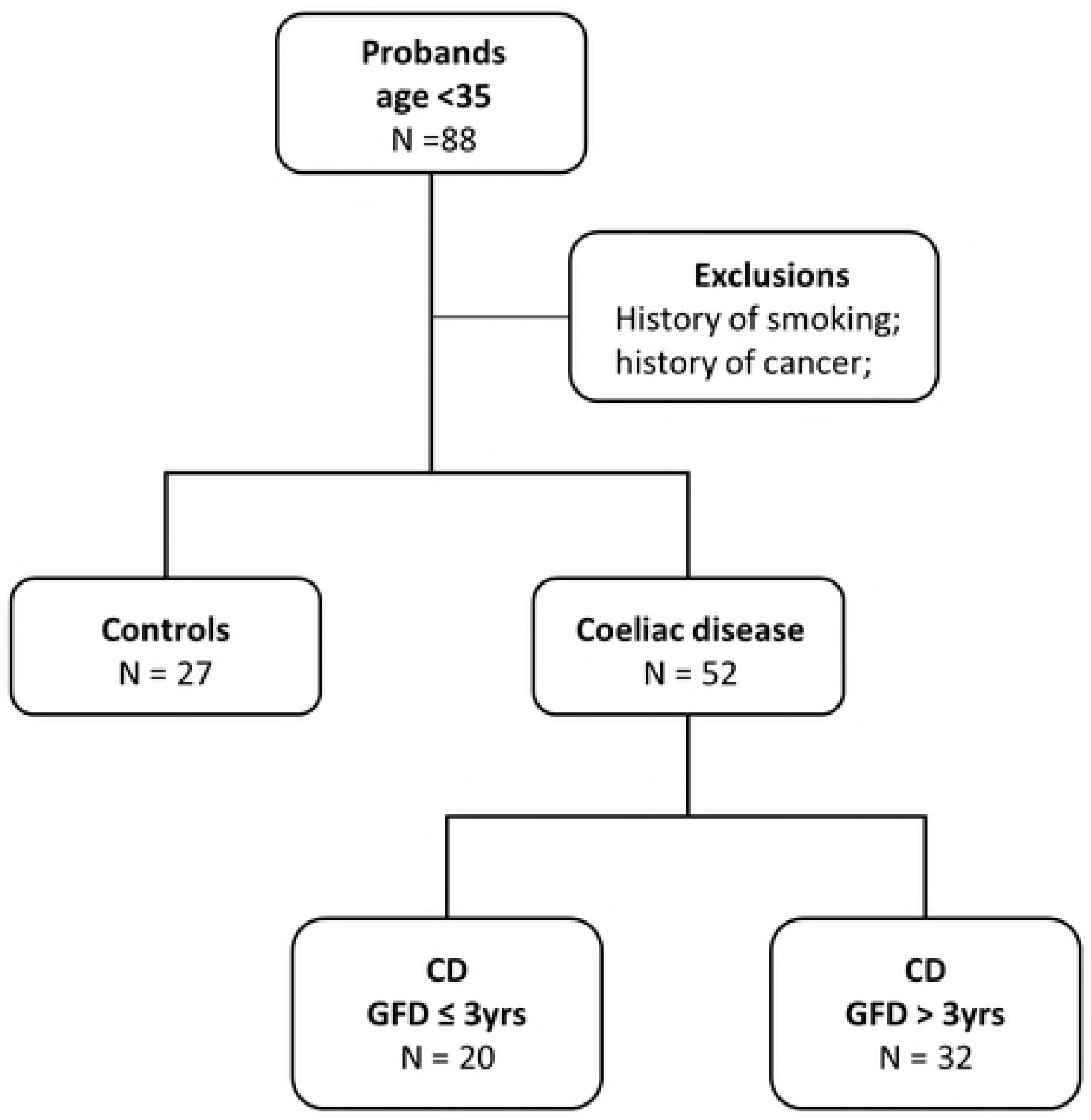
Overview of study participants.

**Figure 2.**
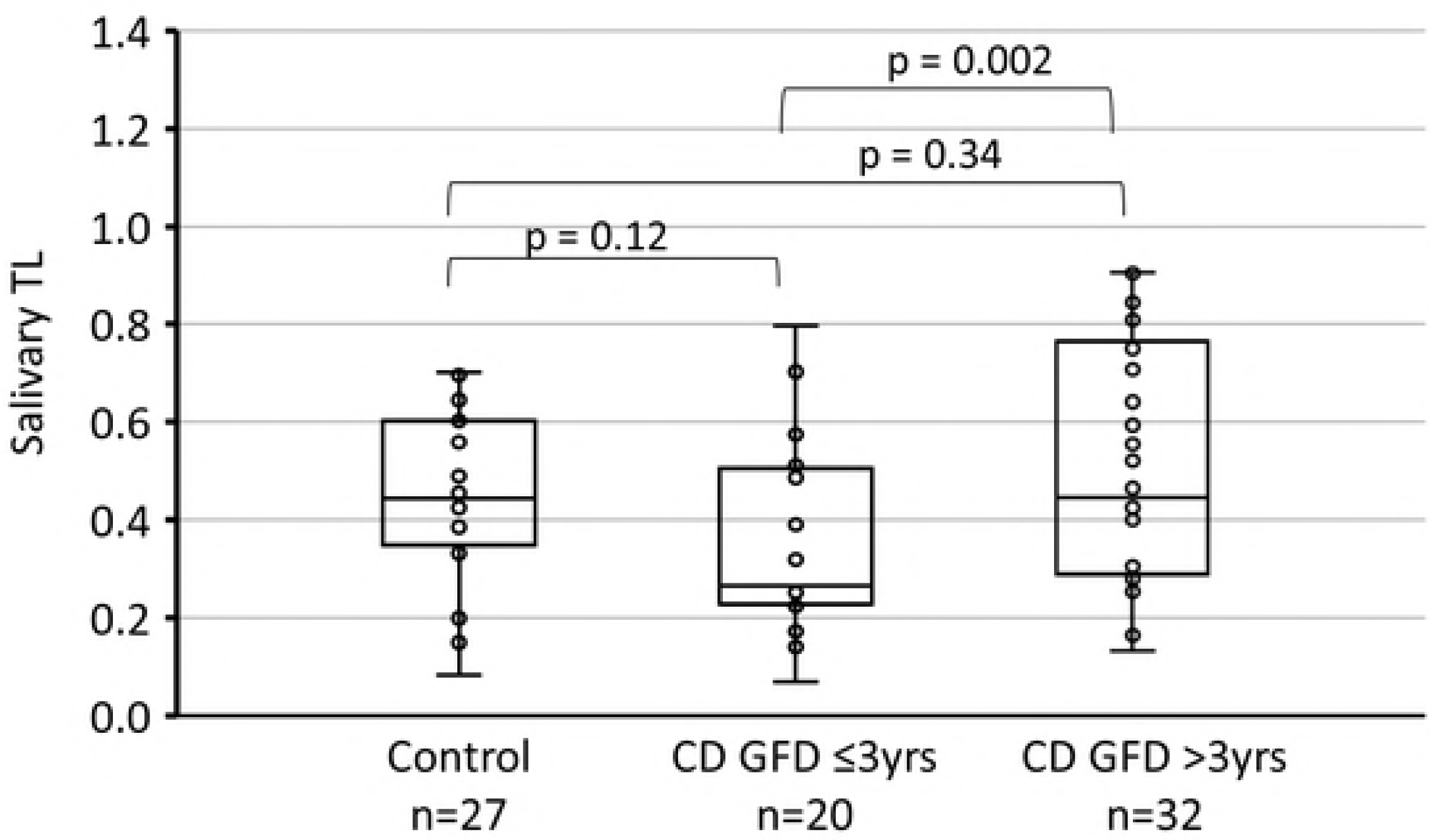
Measurement of salivary telomere length. Salivary telomere length of non-affected controls and individuals with CD who had been on a GFD for ≤ 3 years and >3 years.

In individuals with CD gluten ingestion results in intestinal inflammation and villous atrophy, which can be reversed following removal of dietary gluten [2, 3, 20-22]. Individuals with active CD who are not on a GFD have been reported to have shorter TL when compared to age-matched controls in their intestinal mucosa and peripheral blood leukocytes [1, 13]. The current study was conducted in treated CD individuals who were following a GFD. This may explain why no difference in TL between CD individuals and non-affected controls was observed, as TL is known to recover following the removal of inflammation inducing stressors [9, 23]. Lifestyle interventions including diet that aim to reduce inflammation and oxidative stress have been associated with recovery of TL in individuals recovering from cancer or in high stress professions [24-26]. Similar mechanisms may occur in CD, with TL recovering following reduced inflammation through the removal of dietary gluten. Furthermore, TL was measured in the oral mucosa, not intestinal mucosa or blood, which may have also contributed to the discrepancies observed. The oral mucosa was selected over the other tissue types for its accessibility compared to intestinal mucosa, and because the epithelia and *lamina propria* of oral mucosa reacts to gluten in individuals with CD [27].

For individuals who had been undiagnosed for a long period of time or who had experienced severe disease, the cellular damage to their intestinal mucosa can take many years to completely repair, even with a GFD [28, 29]. This may explain why individuals with CD who had commenced a GFD more recently had significantly shorter TL when compared with individuals who have been treated with a GFD for greater than 3 years. It could also explain why we saw a trend for shorter TL in more recently diagnosed CD individuals when compared to controls, while no difference was observed between CD individuals who had been on GFD for greater than 3 years when compared to controls, as TL also required longer periods of time to recover.

Undiagnosed CD is also associated with an increased risk of developing small intestinal adenocarcinoma and lymphomas [30]. In individuals with CD, this increased risk has been shown to be reduced with a GFD, with CD individuals treated with a GFD for greater than five years having the same risk of cancer as healthy controls [30-32]. It is well established that telomere shortening and telomere dysfunction is a common alteration in the multistep process of malignant transformation in various cancers [33-36]. Telomere attrition may contribute to the increased risk of malignancy in CD [1], however further studies are required to confirm this. Nonetheless, these observations highlight the importance of early diagnosis and treatment of CD to minimise the degree of telomere attrition and risk of malignancy. Early diagnosis and treatment would also enable the repair of intestinal mucosal damage and TL recovery, following the removal of a gluten-induced inflammatory response.

CD has a variable clinical course, some individuals may have had sub-clinical disease for a prolonged period prior to diagnosis, and therefore the years since diagnosis may not always reflect the length of time that the individual had been living with disease. We attempted to reduce this effect by studying a younger population (<35 years of age) who were less likely to have lived with untreated CD for prolonged periods. This study used saliva instead of intestinal mucosa to measure TL, and our findings may not apply to telomeres measured in other tissues of individuals with CD. However, as untreated CD is characterised by systemic inflammation, and the oral mucosa of CD individuals is known to react to gluten exposure, saliva represents an appropriate and non-invasive tissue source to measure TL.

This study measured salivary TL in individuals with treated CD, and non-affected controls. We found no significant difference in salivary TL between individuals with CD and controls. Within the CD cohort, the length of time on a GFD was significantly associated with relative salivary TL, with individuals who had commenced a GFD within the past three years having shorter telomeres compared with individuals who had been treated with a GFD for longer than three years. Treatment of CD with a GFD may assist in the recovery of telomere length, highlighting the importance of early diagnosis and initiation of a GFD to minimise exposure to mucosal damage and telomere attrition.

